# Mitochondrial DNA variation and structure among North American populations of *Megaselia scalaris*

**DOI:** 10.1101/006288

**Authors:** Bret S. Lesavoy, Suzanne E. McGaugh, Mohamed A. F. Noor

## Abstract

The scuttle fly *Megaselia scalaris* is a pest species whose larvae consume living or dead plant or animal tissue, and parasitize humans. Although known to exist on most continents, often transported passively with humans, the connectivity between populations has not been investigated. We use mitochondrial *cytochrome B* sequences to investigate structure among North American isolates of this species. Despite small sample sizes, we detected statistically significant structure among populations. This finding suggests that local measures may be effective in controlling this pest species.

## Introduction

Studying genetic relationships among populations of a species gives insight into patterns of dispersal and gene flow. For pest species, such information is critical for determining the most effective measures of control. For example, if genetic data suggest individuals of a pest species migrate freely among subpopulations, eradication would have to take place in all subpopulations simultaneously to be effective.

The scuttle fly *Megaselia scalaris* is a pest that impacts humans, livestock, and crops. What makes this species unique is the diversity of food on which this omnivorous fly feeds. The scuttle fly feasts on almost any organic or decaying organic material including plants, fungi, and animals (Disney, 2008). *M. scalaris* has also been found breeding in food packets of ship cargoes and has caused economic harm in areas of Texas where the larvae were found in ears of maize (Disney, 2008). Although found worldwide and throughout the United States, it is unclear whether this pest exists as numerous isolated populations or if continental populations experience free and frequent gene flow. If gene flow levels are very high, then local control measures are unlikely to be effective.

The goal of this research project was to determine relatedness of populations (and consequently, degree of gene flow) among isolates of *M. scalaris* collected from across the United States using mitochondrial DNA sequences. Mitochondrial DNA was used because of its ease of primer design, PCR amplification, and sequencing. We attempted to design primers for the NADH2 and srRNA genes to assay the control region, which often exhibits high levels of within-species variation, but we were unable to isolate those sequences successfully. Thus, we focused on the somewhat less variable *cytochrome B* region and analyzed these sequences derived from 12 independent isolates.

## Methods

Strains of *Megaselia scalaris* were obtained via a “citizen science” effort. An advertisement was posted on the *Scientific American* website asking for volunteers. Volunteers were then sent instructions and shipping materials. We identified the subset of samples that were *M. scalaris* and maintained them in culture.

After acquiring *M. scalaris* from different areas of the United States (see Table 1), the Qiagen Gentra Puregene Core Kit A was used to extract DNA from these flies or from one of their descendants. After measuring the DNA yields to confirm that the final isolates had at least 30 ng/ul, we amplified *cytochrome B* via PCR from 1 ul of the genomic DNA. The primers were developed using *M. scalaris* mitochondrial DNA sequences (Rasmussen and Noor, 2009): 5’-

TGAACAAACCTTTACGATCTTCCCATCC-3’ and 5’-

AGGGCGAGCCCCAATTCATGTTAA-3’.

**Table 1:** Number of *Megaselia scalaris* strains analyzed from each collection site.

| State | City | Number of Strains | Donor |
| --- | --- | --- | --- |
| Alabama | Birmingham | 1 | P VanZandt |
| Arizona | Tucson | 1 | M Noor |
| Louisiana | Baton Rouge | 2 | D Donze |
| Michigan | Grand Valley State | 1 | G Sass |
| Nebraska | Lincoln | 3 | S Smith |
| North Carolina | Durham | 2 | S McGaugh / M Noor |
| Texas | Houston | 2 | E Myers-Brunner |

We visualized the PCR products on a 1% TBE agarose gel to confirm that the PCR products properly isolated the expected 1 kb region. We used USB ExoSAP-IT reagent prior to submitting samples for sequencing with Eton Bioscience Inc. The sequence contigs were assembled using Lasergene SeqMan Pro. Once the 12 sequences of the *M. scalaris cytochrome B* region were assembled, we aligned them using BioEdit (Hall, 1999), identified single nucleotide polymorphisms (SNPs), and confirmed them visually in the original electropherograms. We then constructed the phylogeny using MacClade 4.08 (Maddison and Maddison, 2005). Ambiguous indels were excluded from the alignments. The data set was analyzed through a likelihood approach and implemented in GARLI version 2.0 (Zwickl, 2006). Searches were run on the CIPRES computing resource (www.phylo.org; Miller, *et al.*, 2010). Conflict among the resulting phylogeny was assessed according to a 70% maximum likelihood bootstrap (BP) criterion (Mason-Gamer and Kellogg, 1996). In GARLI, a most likely topology was identified and branch support was assessed using a maximum likelihood BS approach. Maximum Likelihood BS analyses (100 replicates) employed a GTR + I + G model of sequence evolution, allowing for estimation of the proportion of invariant sites.

## Results

We sequenced *cytochrome B* in 12 lines of *M. scalaris* from seven populations to identify population structure in North America. We used a maximum likelihood bootstrap tree (100 replicates) to create a majority-rule consensus tree. Using *Episyrphus balteatus* as the outgroup, we found that the 12 lines of *M. scalaris* primarily divided into two clades (Figure 1): one group having strains from Nebraska, North Carolina, and Louisiana, and the other having strains from Michigan, Texas, Arizona, and Alabama. No obvious geographic break was apparent between the sampling locations of the strains representing the two clades. However, none of the four sampling locations from which we obtained multiple strains exhibited both haplotypes.

**Figure 1:**
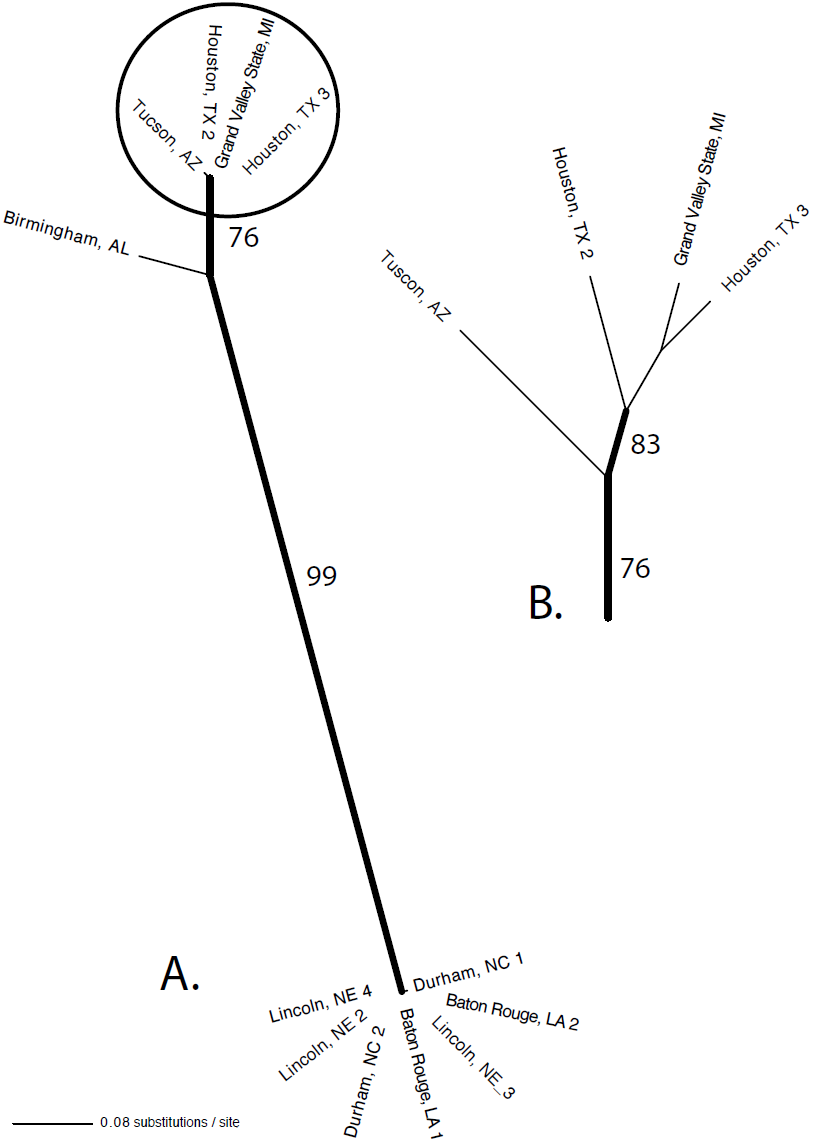
Phylogeny of *Megaselia scalaris* samples. A. Maximum likelihood reconstruction of relationships based on analysis of *cytochrome b* DNA sequences for individuals of *M. scalaris* (-lnL = 1286.2560). Bold branches indicate maximum likelihood bootstrap score > 70% (Mason-Gamer and Kellogg 1996); 100 bootstrap replicates run in Garli 2.0 (Zwickl 2006) on the CIPRES cluster (www.phylo.org; Miller *et al*. 2010). Circled clade expanded in Fig. 1B. B. Cladogram (branch lengths ≠ substitutions / site) based on analyses from Fig. 1A (fully resolved but only bold branches received > 70% bootstrap support).

Although our sample size was small, we explored statistically whether the apparent homogeneity within populations may suggest some structure among or inbreeding within populations. We wrote a short Perl script to resample 12 strains (five bearing one haplotype, seven bearing the other, and having identical numbers of isolates per population as in this study) one million times and identify the number of times all samples from four populations exhibited the same haplotype. Of the one million runs, 23,021 exhibited this pattern, suggesting that if the haplotypes were randomly distributed, we would detect the observed pattern with probability 2.3%. While we failed to detect clear continent-wide structure in our samples, this analysis suggests that North American continental populations of *M. scalaris* are not in panmixia.

## Discussion

We created a phylogeny of mitochondrial sequences from *Megaselia scalaris* using 12 lines from seven North American populations, and observed genetic variation that clustered into two groups (see tree Figure 1). Although no obvious geographic structuring was apparent looking at states of origin, we observed that populations tended to be more homogeneous in haplotype than expected under panmixia.

*M*. *scalaris* is an omnivorous pest that is capable of seeking out and penetrating most organic matter. As a result of its pest-like behavioral characteristics, this species has the potential to spread across the continent through natural dispersal as well as via human transport of organic products. However, its high fecundity and ability to initiate new populations from a single or few gravid females may also cause founder effects, leading to the observed relative mitochondrial gene monomorphism within sampling sites.

Several caveats exist in the interpretation of our results. First, while the *M.* s*calaris* genome contains thousands of genes, we only studied the mitochondrial *cytochrome B* gene. Additionally, the limited variability of the mitochondrial DNA we analyzed prevented us from making more powerful predictions as to the relationships within *M. scalaris* populations. Finally, the small sample size of *M. scalaris* lines we obtained from individual populations limited our scope.

Nonetheless, our demonstration that *M. scalaris* populations are not panmictic suggests that local control measures may be effective in reducing or eliminating particular populations, at least temporarily. Further understanding of the relationships between these distant populations can aid in pest management, and future population genetics studies and can assist in developing *M. scalaris* into a possible model system.

## Acknowledgements

DNA sequences are available in GenBank under accession numbers KJ735473-KJ735484. We thank A. Grusz for performing the phylogenetic analysis and all of the “citizen scientists” who contributed the samples used in our study.

## References

Disney RH. 2008 Natural history of the scuttle fly, Megaselia scalaris. Annu Rev Entomol. 53:39–60.

Hall TA. 1999 BioEdit: a user-friendly biological sequence alignment editor and analysis program for Windows 95/98/NT. Nucl. Acids. Symp. Ser. 41:95–98.

Maddison DR, Maddison WP. 2005. MacClade 4: Analysis of phylogeny and character evolution. Version 4.08a.

Mason-Gamer RJ, Kellogg EA. 1996 Testing for phylogenetic conflict among molecular datasets in the tribe Triticeae (Graminae). Syst. Biol. 45:524–545.

Miller MA, Pfeiffer W, Schwartz T. 2010. Creating the CIPRES science gateway for inference of large phylogenetic trees. In. Proceedings of the Gateway Computing Environments Workshop (GCE); 14 Nov 2010; New Orleans, LA. p. 1–8.

Rasmussen DA, Noor MAF. 2009 What can you do with 0.1x genome coverage? A case study based on a genome survey of the scuttle fly Megaselia scalaris (Phoridae). BMC Genomics. 10:382.

Zwickl D. 2006. Genetic algorithm approaches for the phylogenetic analysis of large biological sequence datasets under the maximum likelihood criterion. [Austin, TX]: University of Texas.

